# Disturbed intramitochondrial phosphatidic acid transport impairs cellular stress signaling

**DOI:** 10.1101/2020.11.05.370098

**Authors:** Akinori Eiyama, Mari J Aaltonen, Hendrik Nolte, Takashi Tatsuta, Thomas Langer

**Author notes:** Corresponding author: Thomas Langer.

## Abstract

Lipid transfer proteins of the Ups1/PRELID1 family facilitate the transport of phospholipids across the intermembrane space of mitochondria in a lipid-specific manner. Heterodimeric complexes of yeast Ups1/Mdm35 or human PRELID1/TRIAP1 shuttle phosphatidic acid (PA) synthesized in the endoplasmic reticulum (ER) to the inner membrane, where it is converted to cardiolipin (CL), the signature phospholipid of mitochondria. Loss of Ups1/PRELID1 proteins impairs the accumulation of CL and broadly affects mitochondrial structure and function. Unexpectedly and unlike yeast cells lacking the cardiolipin synthase Crd1, Ups1 deficient yeast cells exhibit glycolytic growth defects, pointing to functions of Ups1-mediated PA transfer beyond CL synthesis. Here, we show that the disturbed intramitochondrial transport of PA in *ups1*Δ cells leads to altered phospholipid composition of the ER membrane, independent of disturbances in CL synthesis. The impaired flux of PA into mitochondria is associated with the increased synthesis of phosphatidylcholine (PC) and a reduced phosphatidylethanolamine (PE)/PC ratio in the ER of *ups1*Δ cells which suppresses the unfolded protein response (UPR). Moreover, we observed inhibition of TORC1 signaling in these cells. Activation of either UPR by ER protein stress or of TORC1 signaling by disruption of its negative regulator, the SEACIT complex, increased cytosolic protein synthesis and restored glycolytic growth of *ups1*Δ cells. These results demonstrate that PA influx into mitochondria is required to preserve ER membrane homeostasis and that its disturbance is associated with impaired glycolytic growth and cellular stress signaling.

## Introduction

The functional integrity of mitochondria depends on characteristic lipid compositions of their membranes, the mitochondrial outer and inner membrane (OM and IM) (1,2). This is exemplified by the hallmark lipid of mitochondria, cardiolipin (CL), which is pivotal for the structure and function of mitochondria. CL maintains respiration and cristae morphogenesis, ensures protein biogenesis and affects the fusion and fission of mitochondrial membranes (3–6). Altered CL levels in the OM modulate mitophagy and apoptosis (7–11). Hence, reduced levels and aberrant acylation or peroxidation of CL compromise mitochondrial activities and are associated with ageing and various pathophysiological conditions, including cardiomyopathies, skeletal myopathies, ataxias or non-alcoholic fatty liver disease (12–15).

CL is synthesized along an enzymatic cascade at the IM from phosphatidic acid (PA), which is imported from the endoplasmic reticulum (ER), the main cellular phospholipid synthesis center (16). The maintenance of mitochondrial lipid homeostasis requires extensive exchange of phospholipids between the ER and mitochondria (17,18). Similar to other membrane lipids, PA is transported from the ER to the OM at membrane contact sites, which facilitate the bi-directional transport of phospholipids between both organelles. Studies on the cellular distribution of newly synthesized phosphatidylserine (PS) revealed the phospholipid transfer to mitochondria can be driven by localized synthesis (19,20). After reaching the OM, phospholipids are transported across the intermembrane space (IMS) by conserved, lipid-specific lipid transfer proteins (21–26), which likely act in concert with MICOS membrane tethering complexes (22). Heterodimeric complexes of yeast Ups1 (human PRELID1) and Mdm35 (human TRIAP1) serve as PA-specific lipid transfer proteins (21,24). Crystal structures of various members of the conserved Ups1/PRELID1 family of lipid transfer proteins show an internal lipid binding pocket, which is surrounded by a β-sheet and α-helices, reminiscent of other classes of lipid transfer proteins (27–30). Loss of Ups1 or PRELID1 in yeast or human cells, respectively, inhibits intramitochondrial PA transport and limits CL synthesis at the IM, which is accompanied by mitochondrial deficiencies, such as impaired respiration and mitochondrial fragmentation, and increases the susceptibility of the cells towards apoptotic stimuli (21,24,31–33).

High levels of PRELID1 and TRIAP1 were identified as an unfavorable prognostic marker in cancer (34). As cancer cells are often glycolytic, it is noteworthy that yeast cells lacking Ups1 also exhibit slow growth in glucose media (21,32,33). This phenotype is difficult to reconcile with reduced CL levels only, as cells lacking the cardiolipin synthase Crd1 in mitochondria grow normally in glucose-containing media although they are virtually devoid of CL (35). It therefore appears that the inhibition of intramitochondrial PA transport upon loss of Ups1 does not only perturb the mitochondrial membrane homeostasis, but also causes deficiencies presumably beyond mitochondrial functions. Here, we demonstrate that a disturbed intramitochondrial transport of PA affects TORC1 signaling and alters the phospholipid homeostasis of the ER membrane modulating the unfolded protein response (UPR).

## Results

### The loss of Ups1 impairs basal UPR

To examine how the loss of Ups1 affects cell growth under glycolytic conditions, we used quantitative mass spectrometry (qMS) to determine the phospholipid profile of cellular membranes in wild-type and *ups1*Δ cells which were grown in glucose-containing media. In agreement with previous findings (21,32), we observed reduced CL levels in *ups1*Δ cells reflecting impaired PA transport to CL-synthesizing enzymes at the IM (Fig. 1, A and B, and Fig. S1A). We also noted a significant increase of phosphatidylcholine (PC) and a decrease of phosphatidylethanolamine (PE), resulting in a low PE/PC ratio in cellular membranes (Fig. 1, A to C, and Fig. S1A). Unexpectedly, deletion of *UPS1* also affected the phospholipid composition of ER membranes. We observed an increased total phospholipid content of an ER-enriched membrane fraction, which was accompanied by a significant increase of PC and phosphatidylinositol (PI) (Fig. 1, D and E, and Fig. S1B), and a low PE/PC ratio (Fig. 1F).

**Figure 1.**
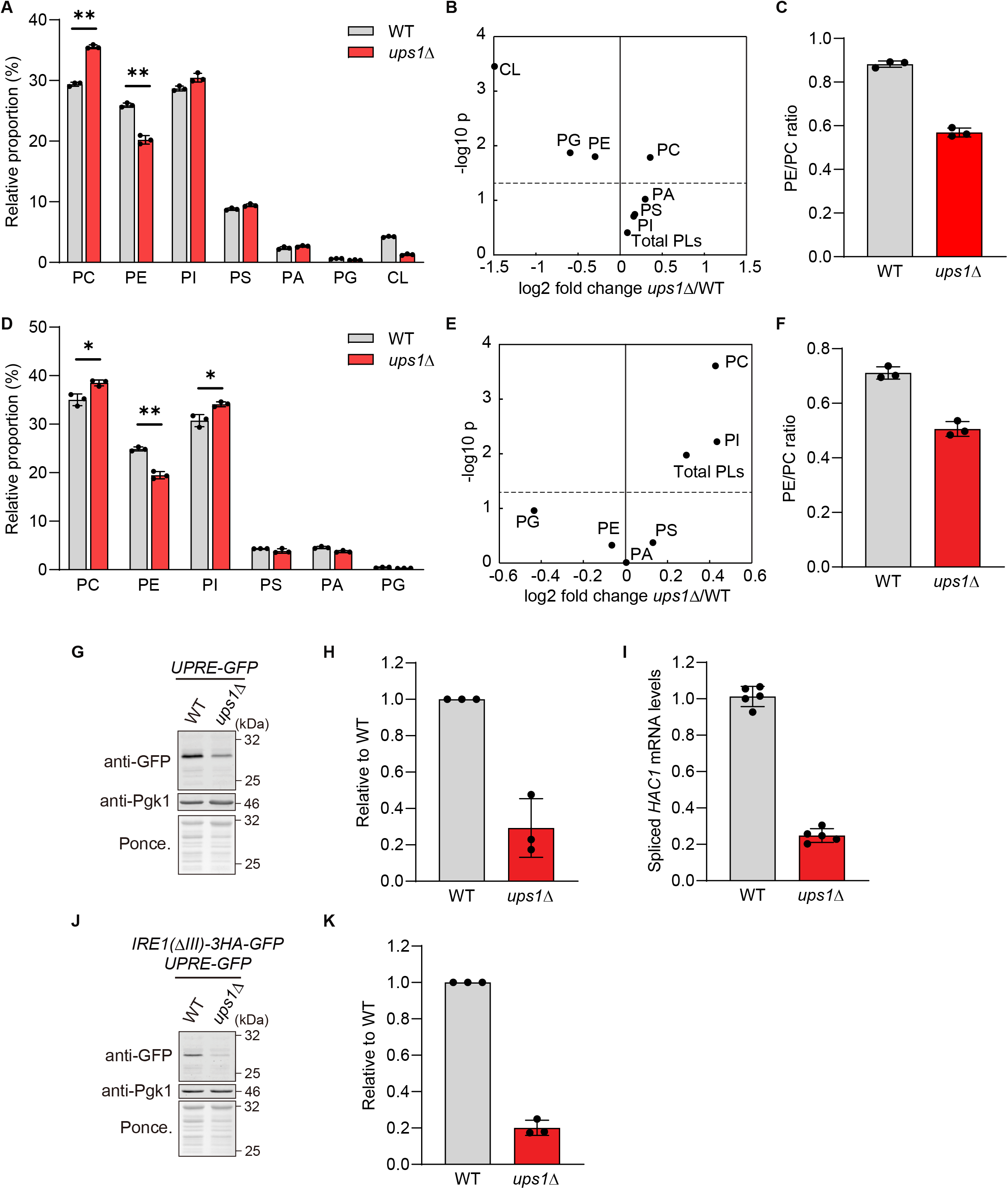
ER phospholipid composition is altered in *ups1*Δ cells. (A) Phospholipid composition in whole cell of wild-type (WT) and *ups1*Δ cells grown to log phase in SCD medium. Data represent mean ± SD (n = 3). \*\**P* < 0.01 (B) Changes in the absolute abundance of phospholipids in whole cell of *ups1*Δ. Data represented as log2 fold change at x-axis with -log 10 p value of student’s t-test at y-axis. Dashed line represents *p* = 0.05. (C) PE/PC ratio of WT and *ups1*Δ cells. (D) Phospholipid composition in ER-enriched microsome fraction from WT and *ups1*Δ cells grown to log phase in SCD medium. Data represent mean ± SD (n = 3). \**P* < 0.05, \*\**P* < 0.01. (E) Changes in the absolute abundance of phospholipids in ER-enriched microsome fraction of *ups1*Δ. Data represented as log2 fold change at x-axis with -log 10 p value of student’s t-test at y-axis. Dashed line represents *p* = 0.05. (F) PE/PC ratio in ER-enriched microsome fraction from WT and *ups1*Δ cells. (G) WT and *ups1*Δ cells expressing *4xUPRE-GFP* grown to log phase in SCD medium were subjected to western blotting. Pgk1 and ponceau staining were monitored as a loading control. (H) GFP in (G) was quantified. GFP signals were normalized to Pgk1 and expressed relative to WT cells (set as one). Data represent mean ± SD (n = 3). (I) WT and *ups1*Δ cells were grown to log phase in SCD medium. Spliced *HAC1* mRNA expression was analyzed by real time PCR and normalized to *ACT1* mRNA expression. Data represent mean ± SD (n = 5). (J) WT and *ups1*Δ cells expressing Ire1(ΔIII)-3HA-GFP and *4xUPRE-GFP* grown to log phase in SCD medium were subjected to western blotting. (K) GFP in (J) was quantified as described in (H). Data represent mean ± SD (n = 3).

Previous studies have reported that an increased PE/PC ratio in ER membranes activates the unfolded protein response (UPR) independent of protein stress (36–39). We therefore monitored UPR induction in *ups1*Δ cells harboring GFP under the control of the UPR element (UPRE) in their genome (40). Whereas GFP accumulated in wild-type cells under basal conditions, GFP expression was decreased in cells lacking Ups1 (Figure 1, G and H). Moreover, we observed reduced mRNA levels of spliced *HAC1*, encoding a transcription factor involved in UPR (Figure 1I) (41–43). We therefore conclude that basal gene expression under the control of UPRE is suppressed in *ups1*Δ cells. Notably and in contrast to *ups1*Δ cells, cells lacking other enzymes involved in CL synthesis (such as Tam41, Pgs1 or Crd1) did not show significant reduction of the GFP expression under UPRE (Fig. S1, C to F), pointing that CL defect per se does not lead to the suppression of UPR.

A central sensor of UPR is the ER kinase Ire1, which senses unfolded proteins in the ER lumen as well as lipid bilayer stress in the membrane (41,44–46). To assess the contribution of lipid bilayer stress specifically, we exploited the previously described ΔIII mutant of Ire1, which is able to sense lipid bilayer stress but not unfolded ER proteins (47,48). Cells expressing Ire1(ΔIII) recapitulated the suppression of the UPR by the loss of Ups1 (Fig. 1, J and K). These results suggest that the altered phospholipid composition of ER membranes limits UPR activation in *ups1*Δ cells.

### Increased PC levels cause suppression of basal UPR

To examine directly how a decreased PE/PC ratio in ER membranes affects UPR, we induced *de novo* synthesis of PC via the Kennedy pathway by supplementing the growth medium with choline (49) and monitored UPR activity. Choline supplementation increased PC levels in ER-enriched membrane fractions of wild-type cells and led to a lower PE/PC ratio in these membranes (Fig. 2, A and B). Similar to cells lacking Ups1, the decreased PE/PC ratio in ER membranes was accompanied by an impaired basal UPR (Fig. 2, C and D), which was not altered in cells expressing mutant Ire1(ΔIII) (Fig. 2, E and F). We conclude from these experiments that the basal UPR is impaired if the PE/PC ratio in ER membranes is decreased, whereas an increased PE/PC ratio in ER membranes induces the UPR (39).

**Figure 2.**
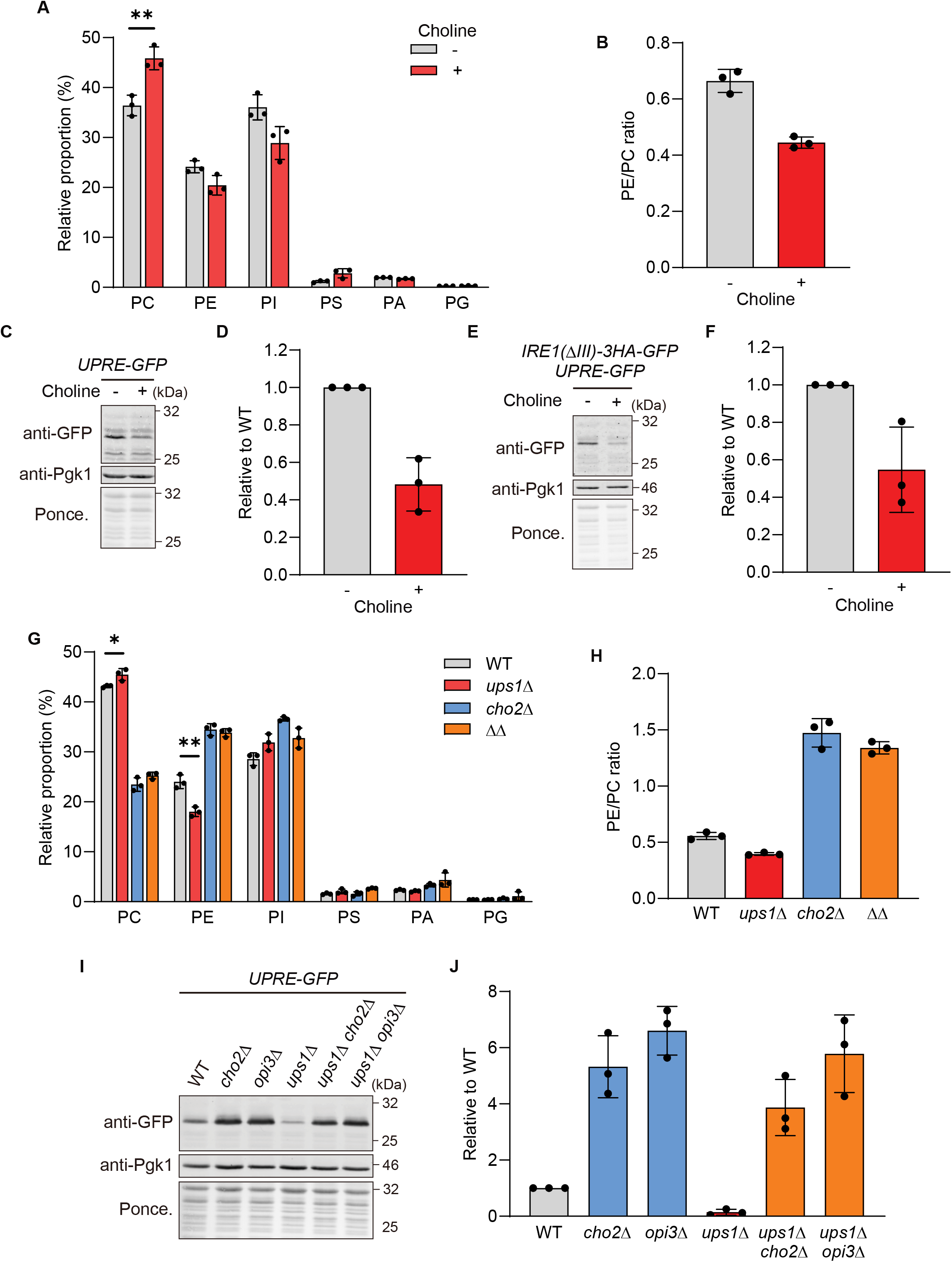
Basal UPR is suppressed in high PC levels. (A) Phospholipid composition in ER-enriched microsome fraction from wild-type (WT) cells grown to log phase in SCD medium with or without 1 mM choline. Data represent mean ± SD (n = 3). (B) PE/PC ratio in ER-enriched microsome fraction from WT with or without choline. (C) WT cells expressing *4xUPRE-GFP* grown to log phase in SCD medium with or without 1 mM choline were subjected to western blotting. Pgk1 and ponceau staining were monitored as a loading control. (D) GFP in (C) was quantified. GFP signals were normalized to Pgk1 and expressed relative to WT cells (set as one). Data represent mean ± SD (n = 3). (E) WT cells expressing Ire1(ΔIII)-3HA-GFP and *4xUPRE-GFP* grown to log phase in SCD medium with or without 1 mM choline were subjected to western blotting. (F) GFP in (E) was quantified as described in (D). Data represent mean ± SD (n = 3). (G) Phospholipid composition in ER-enriched microsome fraction from WT, *ups1*Δ, *cho2*Δ and *ups1*Δ*cho2*Δ (ΔΔ) cells grown to log phase in SCD medium. Data represent mean ± SD (n = 3). \**P* < 0.05, \*\**P* < 0.01. (H) PE/PC ratio of ER-enriched microsome fraction from WT, *ups1*Δ, *cho2*Δ and *ups1*Δ*cho2*Δ (ΔΔ) cells. (I) WT, *cho2*Δ, *opi3*Δ, *ups1*Δ, *ups1*Δ*cho2*Δ, and *ups1*Δ*opi3*Δ cells expressing *4xUPRE-GFP* grown to log phase in SCD medium were subjected to western blotting. (J) GFP in (I) was quantified as described in (D). Data represent mean ± SD (n = 3).

To unambiguously demonstrate that Ups1 affects UPR activation by modulating the phospholipid profile in ER membranes, we manipulated the PE/PC ratio in *ups1*Δ cells genetically. Yeast cells lacking Cho2 or Opi3, which are required for synthesis of PC from PE (49), accumulate PE relative to PC (Fig. 2G). Accordingly, deletion of *CHO2* or *OPI3* induces the UPR (39). We therefore deleted *CHO2* or *OPI3* in *ups1*Δ cells, which strongly increased the PE/PC ratio in an ER-enriched membrane fraction (Fig. 2, G and H). This was accompanied by UPR activation in the absence of Ups1 (Fig. 2, I and J). Thus, *ups1*Δ cells maintain the ability to induce the UPR upon lipid bilayer stress, but the reduced PE/PC ratio in the ER membrane of *ups1*Δ cells impairs the UPR under basal conditions.

### UPR activation restores glycolytic growth of *ups1*Δ cells

Having established the link between Ups1 and UPR activation, we examined in further experiments whether the reduced UPR activity can explain the impaired glycolytic growth of *ups1*Δ cells. We therefore treated *ups1*Δ cells with tunicamycin, an inhibitor of protein glycosylation, which causes ER protein stress and induces the UPR. Treatment with tunicamycin activated the UPR both in wild-type and in *ups1*Δ cells (Fig. 3A) and allowed growth of *ups1*Δ cells on fermentable medium (Fig. 3B). The restoration of cell growth by tunicamycin depended on Ire1-Hac1 signaling (Fig. 3C), demonstrating that UPR activation is sufficient to promote glycolytic growth of *ups1*Δ cells. Consistently, activation of the UPR upon overexpression of Ire1 (Fig. 3D) (45,50) or deletion of *CHO2* or *OPI3* (Fig. 2, I and J) suppressed growth deficiencies of *ups1*Δ cells on glucose-containing medium (Fig. 3, E and F). Notably, previous studies correlated the growth of *ups1*Δ cells on glucose-containing media with mitochondrial CL levels (51). However, overexpression of Ire1 did not alter the accumulation of CL (Fig. 3G). We therefore conclude that UPR activation is sufficient to allow growth of *ups1*Δ cells on fermentable carbon sources independent of CL levels in mitochondrial membranes.

**Figure 3.**
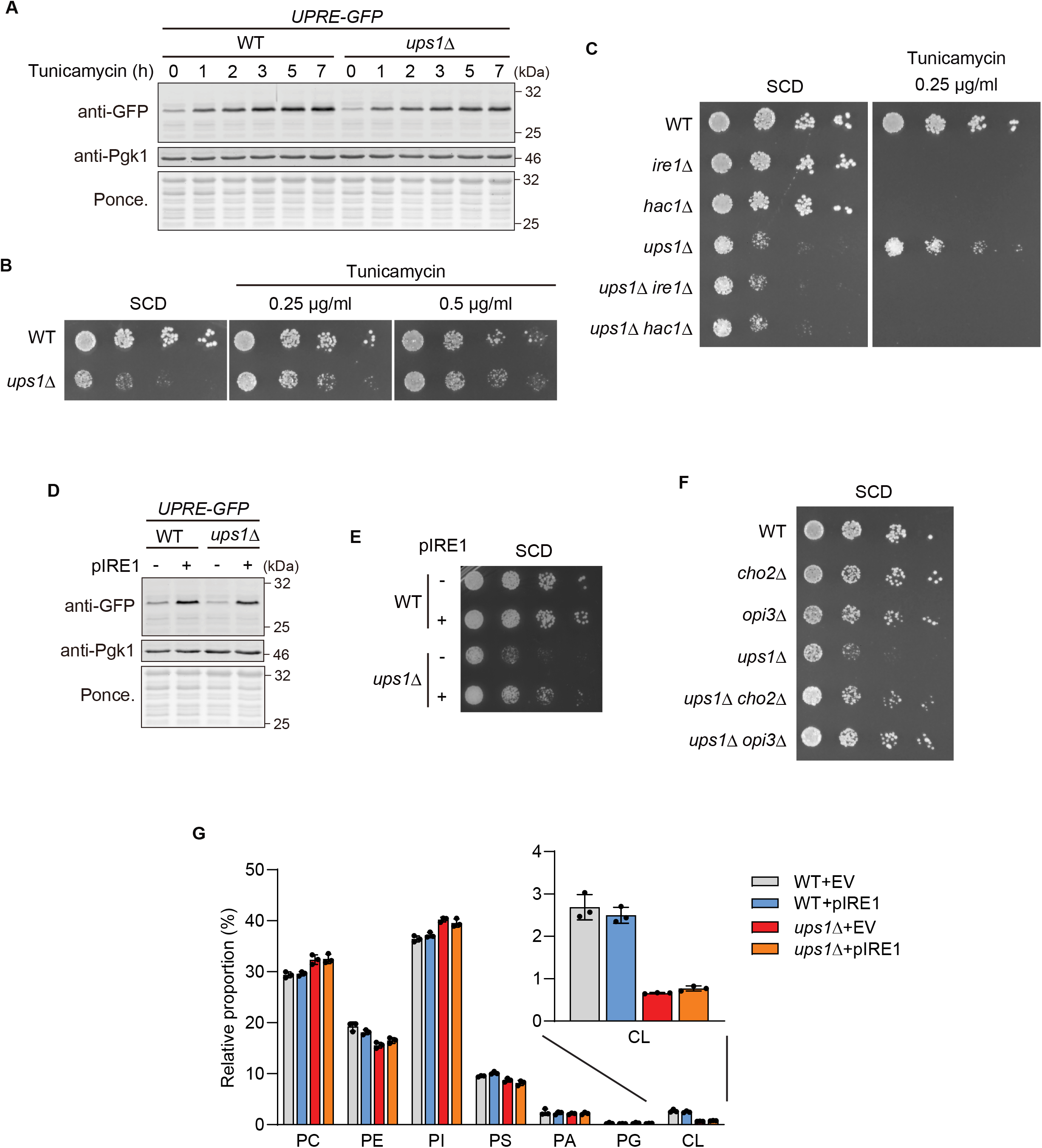
Glycolytic growth of *ups1*Δ cells is restored upon UPR activation. (A) Wild-type (WT) and *ups1*Δ cells expressing *4xUPRE-GFP* grown to log phase in SCD medium were treated with 1 μg/ml tunicamycin, collected at the indicated time points and subjected to western blotting. Pgk1 and ponceau staining were monitored as a loading control. (B) Serial dilutions of WT and *ups1*Δ cells were spotted on SCD medium with or without tunicamycin (concentration as 0.25 or 0.5 μg/ml) and incubated at 30°C for 2 day (n = at least 3). (C) Serial dilutions of WT, *ire1*Δ, *hac1*Δ, *ups1*Δ, *ups1*Δ*ire1*Δ, *ups1*Δ*hac1*Δ cells were spotted on SCD medium with or without 0.25 μg/ml tunicamycin and incubated at 30°C for 2 day (n = at least 3). (D) *4xUPRE-GFP* expressing WT and *ups1*Δ cells transformed with an empty vector or a plasmid encoding Ire1 (pIRE1) were grown to log phase in SCD medium and subjected to western blotting. (E) Serial dilutions of WT and *ups1*Δ cells transformed with an empty vector or pIRE1 were spotted on SCD medium and incubated at 30°C for 2 days (n = at least 3). (F) Serial dilutions of WT, *cho2*Δ, *opi3*Δ, *ups1*Δ, *ups1*Δ*cho2*Δ, and *ups1*Δ*opi3*Δ cells were spotted on SCD medium and incubated at 30°C for 2 days (n = at least 3). (G) Phospholipid composition in whole cell of WT and *ups1*Δ cells transformed with an empty vector (EV) or pIRE1. Data represent mean ± SD (n = 3).

### Reduced cellular protein synthesis in *ups1*Δ cells

We noted that the loss of UPR activity in *ire1Δ* and *hac1Δ* cells did not limit growth in glucose-containing medium (Fig. 3C). This contrasts *ups1*Δ cells and therefore points to additional deficiencies in *ups1*Δ cells that limit cell growth. We therefore determined transcriptome and proteome profiles of *ups1*Δ cells under glycolytic growth conditions (Table S1 and S2). Consistent with the observed impaired UPR activity, canonical UPR target genes were expressed at lower levels in *ups1*Δ cells when compared to wild-type cells (Fig. S2A). Both transcriptome and proteome profiles pointed to reduced protein synthesis in *ups1*Δ cells (Fig. 4, A and B). The expression of genes involved in processes driving protein synthesis, such as rRNA processing or ribosome biogenesis, was most prominently downregulated in *ups1*Δ cells (Fig. 4A and Table S3). Consistently, proteins regulating rRNA processing accumulated at reduced levels in *ups1*Δ cells (Fig 4B and Table S4). Together, these data demonstrate that protein synthesis is inhibited in Ups1-deficient cells. The expression of genes related to protein synthesis is decreased under stress conditions (52). Consistently, tunicamycin treatment limited the expression of genes involved in cellular protein synthesis (Fig. 4C). However, it partially promoted cellular protein synthesis in *ups1*Δ cells (Fig. 4D), consistent with the improved cell growth under these conditions (Fig. 3B).

**Figure 4.**
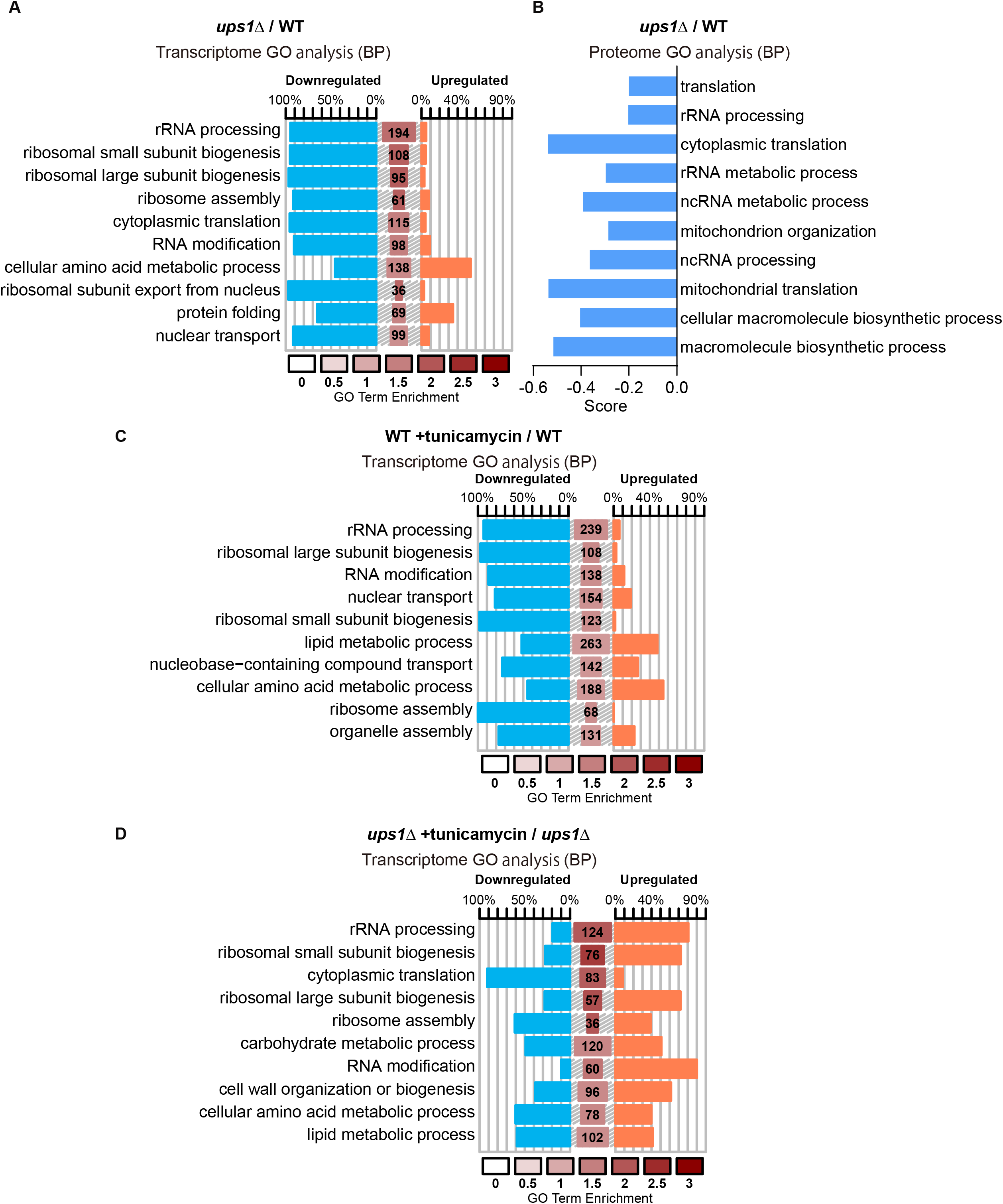
*ups1*Δ cells show suppressed ribosome synthesis. (A) Symplot representing the changed genes within an enriched biological process (BP) GO terms in *ups1*Δ cells against wild-type (WT) cells. Top 10 significant terms from Yeast GO-slim are plotted. Percentage of down (blue) and up (red) regulated genes in each term is shown. The size of the central bars was directly proportional to the number of genes in the query belonging to the respective term and the enrichment value is represented by the color of the bar as previously reported (67). (B) 1D annotation enrichment analysis of proteome in *ups1*Δ cells against WT cells is shown (71). Annotation is biological process (BP) GO terms. Score represents difference of the distribution of the ratios for the proteins corresponding to GO terms from the ratios of the distribution for all proteins. Top 10 significant terms from Yeast GO-slim are indicated. (C) Symplot representing the changed genes within an enriched biological process (BP) GO terms in WT cells treated with 1 μg/ml tunicamycin for 3 h against non-treated WT cells. Top 10 significant terms from Yeast GO-slim are plotted. (D) Symplot representing the changed genes within an enriched biological process (BP) GO terms in *ups1*Δ cells treated with 1 μg/ml tunicamycin for 3 h against non-treated *ups1*Δ cells. Top 10 significant terms from Yeast GO-slim are plotted.

### TORC1 inhibition limits glycolytic growth of *ups1*Δ cells

The target of rapamycin complex 1 (TORC1) regulates ribosome biogenesis in response to the metabolic status of cells (53). We noted an increased expression of genes coding core autophagy proteins in *ups1*Δ cells when compared to wild-type cells (Fig. S2B and Table S1), suggesting inhibition of TORC1 and induction of autophagy in *ups1*Δ cells. To directly assess TORC1 activity in *ups1*Δ cells, we monitored the level and phosphorylation of one of its substrates, Sch9 (54), an AGC family protein kinase, which is homologous to mammalian S6 and Akt kinases and regulates ribosome biogenesis (55,56). We found that Sch9 protein levels were not changed (Fig. 5, A and B), but Sch9 phosphorylation was significantly suppressed in *ups1*Δ cells compared to wild-type cells (Fig. 5, C and D). These results indicate low TORC1 activity in *ups1*Δ cells, which agrees with the reduced protein synthesis in these cells.

**Figure 5.**
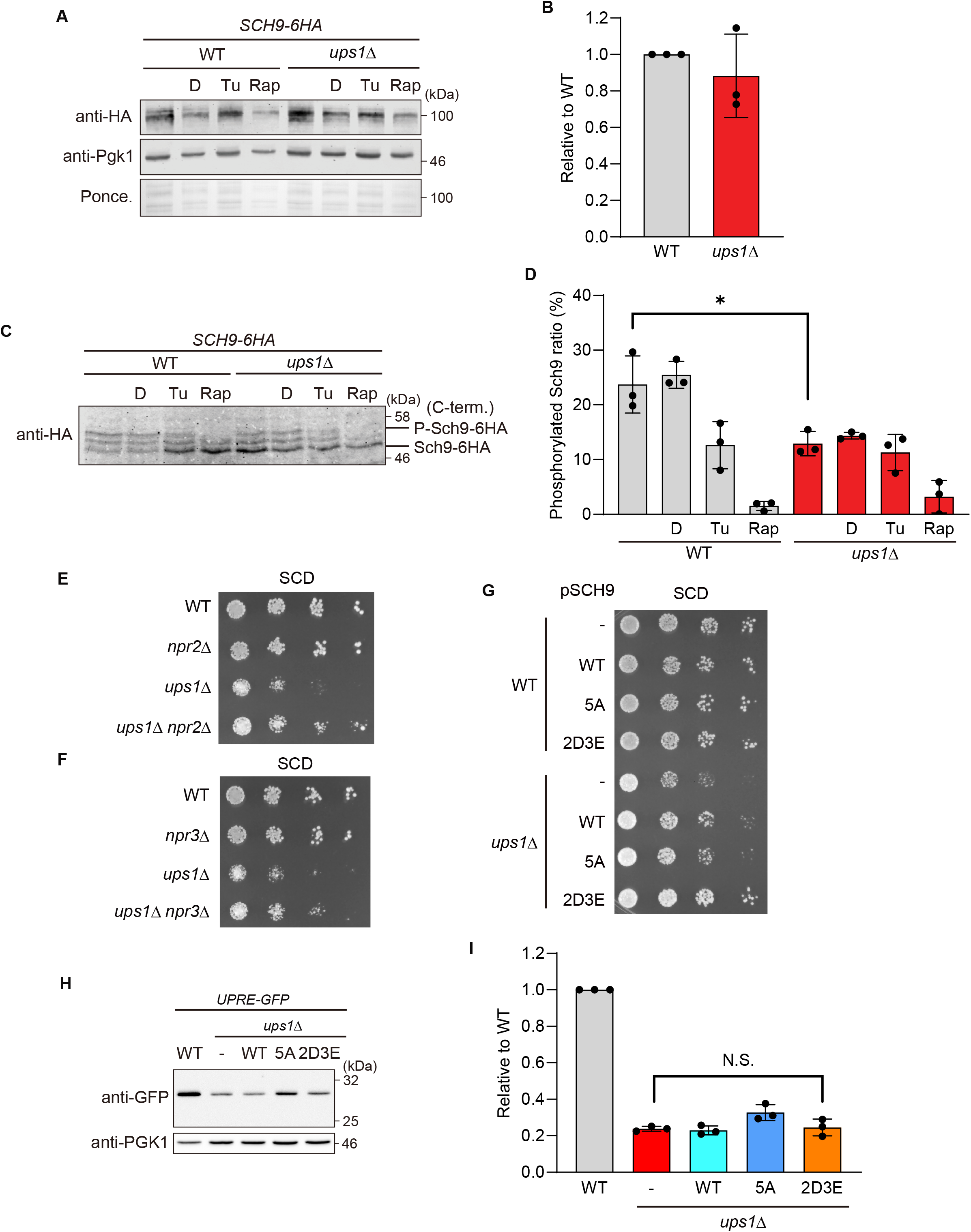
TORC1-Sch9 signaling is impaired in *ups1*Δ cells. (A) Wild-type (WT) and *ups1*Δ cells expressing Sch9-6HA grown to log phase in SCD medium were treated with DMSO (D), 1 μg/ml tunicamycin (Tu), or 0.2 μg/ml rapamycin (Rap) for 3 h and subjected to western blotting. Pgk1 and ponceau staining were monitored as a loading control. (B) Sch9-6HA of non-treated cells in (A) was quantified. Sch9-6HA signals were normalized to Pgk1 and expressed relative to wild-type cells (set as one). Data are means ± SD (n = 3). (C) WT and *ups1*Δ cells expressing Sch9-6HA grown to log phase in SCD medium were treated with DMSO (D), 1 μg/ml tunicmycin (Tu), or 0.2 μg/ml rapamycin (Rap) for 3 h. For the analysis of Sch9 phosphorylation, lysates were treated with NTCB and subjected to western blotting. (D) Phosphorylated Sch9-6HA ratio in (C) was quantified. Phosphorylated Sch9-6HA signals were divided with total Sch9-6HA signals. Data represent mean ± SD (n = 3). \**P* < 0.05. (E) Serial dilutions of WT, *npr2*Δ, *ups1*Δ, and *ups1*Δ*npr2*Δ cells were spotted on SCD medium and incubated at 30°C for 2 days (n = 3). (F) Serial dilutions of WT, *npr3*Δ, *ups1*Δ, and *ups1*Δ*npr3*Δ cells were spotted on SCD medium and incubated at 30°C for 2 days (n = 3). (G) Serial dilutions of WT and *ups1*Δ cells transformed with an empty vector or a plasmid encoding Sch9^WT^ (pSCH9), Sch9^5A^ (T723A, S726A, T737A, S758A, S765A), or Sch9^2D3E^ (T723D, S726D, T737E, S758E, S765E) were spotted on SCD medium and incubated at 30°C for 2 days (n = at least 3). (H) *4xUPRE-GFP* expressing WT and *ups1*Δ cells transformed with an empty vector or a plasmid encoding Sch9^WT^, Sch9^5A^, or Sch9^2D3E^ were grown to log phase in SCD medium and subjected to western blotting. (I) GFP in (H) was quantified. GFP signals were normalized to Pgk1 and expressed relative to WT cells (set as one). Data represent mean ± SD (n = 3). N.S., not significant.

We hypothesized that, similar to UPR activity, an altered membrane lipid composition may inhibit TORC1 activity in *ups1*Δ cells. However, neither loss of the CL synthase Crd1 to decrease CL levels (Fig. S3A), nor choline supplementation of wild-type cells to mimic the PE/PC ratio of *ups1*Δ cells affected TORC1 activity and phosphorylation of Sch9 (Fig. S3, B to E). Similarly, decreased PE levels upon loss of the mitochondrial phosphatidylserine (PS) transfer protein Ups2 did not alter TORC1 activity (Fig. S3, B and C). TORC1 activity is dependent on PA (57,58), but disturbed intramitochondrial PA transport in *ups1*Δ cells did not result in the accumulation of PA in ER-enriched membrane fractions (Fig. 1, D and E). Moreover, PA accumulation upon inhibition of the PA phosphatase in *nem1Δ* or *spo7Δ* cells did not affect Sch9 phosphorylation (Fig. S3, F and G). Thus, other deficiencies appear to inhibit TORC1 activity in *ups1*Δ cells.

To examine whether TORC1 inhibition limits the glycolytic growth of *ups1*Δ cells, we used a genetic approach. The Seh1-associated complex inhibiting TORC1 (SEACIT) consisting of Npr2, Npr3 and Iml1 functions as negative regulator of TORC1 in yeast (59,60). Constitutive activation of TORC1 upon deletion of one of the SEACIT subunits restored cell growth of *ups1*Δ cells (Fig. 5, E and F). Moreover, expression of a phosphomimetic mutant variant of Sch9, Sch9^2D3E^, greatly restored the growth of *ups1*Δ (Fig. 5G). We therefore conclude that inhibition of TORC1-Sch9 signaling impairs the growth of *ups1*Δ cells on a fermentable carbon source such as glucose. Notably, TORC1 activation upon expression of Sch9^2D3E^ in *ups1*Δ cells did not restore UPR activity (Fig. 5, H and I), indicating that both pathways promote glycolytic growth of *ups1*Δ cells independently.

## Discussion

Ups1-dependent transport of PA to the IM promotes CL synthesis (21). Here, we have unraveled an unexpected link between intramitochondrial PA transport and cellular signaling (Fig. 6). Disturbed PA transport in mitochondria lacking Ups1 limits the activation of UPR and inhibits TORC1. TORC1 signaling controls ribosome biogenesis and protein synthesis, which is suppressed in *ups1*Δ cells and limits glycolytic cell growth. Accordingly, activation of either UPR or TORC1 is sufficient to suppress growth deficiencies in the absence of Ups1. Thus, the flux of PA into mitochondria does not only ensure mitochondrial CL synthesis and membrane biogenesis, but is coupled to the phospholipid homeostasis of other cellular membranes affecting cell signaling.

**Figure 6.**
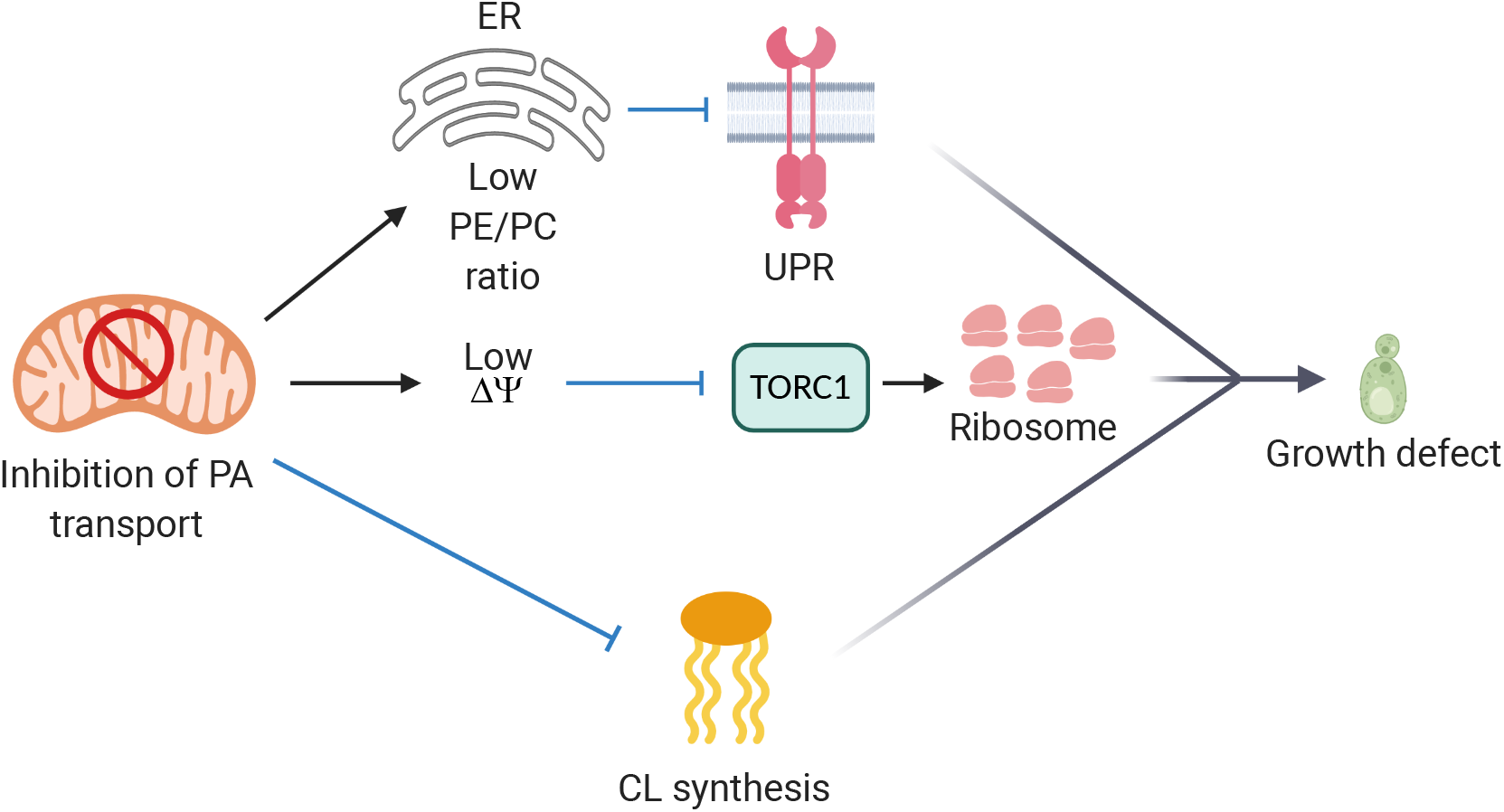
Altered PA transport in mitochondria affects UPR activation and TORC1 signaling. Impaired PA transport in mitochondria upon loss of Ups1 alters the ER lipid composition (low PE/PC ratio) inhibiting UPR. Independently, TORC1-Sch9 signaling is suppressed in *ups1*Δ cells. Deficiencies in UPR and TORC1 signaling combined with reduced CL synthesis limit cell growth.

Several lines of evidences reveal that the disturbed intramitochondrial PA transport in *ups1*Δ cells impairs independently CL synthesis, UPR and TORC1 signaling. The activation of the UPR allowed glycolytic growth of *ups1*Δ cells, but did not restore the accumulation of CL in mitochondrial membranes (Fig. 3G) or the phosphorylation of Sch9 (Fig. 5, C and D). On the other hand, mimicking the PE/PC ratio observed in *ups1*Δ cells by choline supplementation of wild type cells suppressed UPR, but did not alter the phosphorylation of Sch9 (Fig. S3, D and E). Conversely, expression of phosphomimetic Sch9^2D3E^ mutant did not restore basal UPR in *ups1*Δ cells (Fig. 5, H and I). These findings are consistent with a previous study demonstrating that activation of TORC1 does not affect the UPR (61). Moreover, CL deficiencies in cells lacking the cardiolipin synthase Crd1 did not suppress TORC1 signaling (Fig. S3, B and C) or basal UPR (Fig. S1, E and F). Therefore, we conclude that the impaired intramitochondrial PA transport rather than the CL deficiency leads to the alteration of stress signaling cascades outside of *ups1*Δ mitochondria.

Inhibition of PA transport across the IMS in *ups1*Δ cells is associated with an increased PC and PI and a decreased PE/PC ratio but not with a significant accumulation of PA in the ER membrane. This indicates efficient bi-directional transport of PA between the OM and the ER membrane and suggests that PA is rapidly metabolized in the ER, if it is not consumed for CL synthesis in *ups1*Δ mitochondria. Since PA serves as a common precursor for the synthesis of PC and CL, PA is likely converted to PC resulting in lowered PE/PC ratios in the ER membrane and the inhibition of UPR. Rapid conversion to PC can also explain why impaired PA utilization in mitochondria does not suppress Opi1, the transcriptional repressor of phospholipid synthesis, which is regulated by PA (49). Transcript profiling revealed no increased expression of Opi1 target genes in *ups1*Δ cells, consistent with the unaltered PA levels and increased PI in ER membranes in *ups1*Δ cells. The uncontrolled accumulation of PC and PI in the ER membrane may also retard functionality of the ER in addition to the limited UPR activation, contributing to the impaired glycolytic growth in *ups1*Δ cells.

The low PE/PC ratio in ER membranes suppresses the basal UPR in *ups1*Δ cells, complementing previous findings which revealed UPR activation upon an increase of the PE/PC ratio (Thibault et al., 2012). The ratio of PE and PC determines membrane packing and curvature stress in the ER membrane, which appears to be sensed by the Ire1 kinase. Indeed, an amphipathic helix adjacent to the transmembrane domain of Ire1 was found to detect biophysical changes at the ER membrane (62). It therefore can be envisioned that the low PE/PC ratio may improve membrane packing and restrain the formation of the X-shaped transmembrane dimer of Ire1.

Protein synthesis is suppressed under stress conditions (52). Our transcriptome data correspondingly show that expression levels of genes involved in ribosome biogenesis are decreased in wild-type cells treated with tunicamycin. On the other hand, tunicamycin induces a number of those genes in *ups1*Δ cells in accordance with the recovery of glycolytic growth of the cells. One remaining question is how tunicamycin treatment induces genes involved in protein synthesis, although it does not restore TORC1 signaling in *ups1*Δ cells. In contrast to the previous indication that the suppression of protein synthesis with tunicamycin proceeds independent of the UPR (63), the growth recovery of *ups1*Δ cells by tunicamycin depends on Ire1 and Hac1. Therefore, our results imply the existence of an UPR-dependent pathway promoting protein synthesis that is induced by proteotoxic stress in *ups1*Δ cells.

The loss of Ups1 also impairs TORC1 signaling independent of UPR inhibition. TORC1 inhibition in *ups1*Δ cells is not caused by alterations in the membrane lipid composition of the ER membrane, as neither a decreased PE/PC ratio, nor the accumulation of PA by other means (e.g. inhibition of PA phosphatase complex in *nem1Δ* or *spo7Δ* cells (Fig. S3, F and G)) resulted in the inhibition of TORC1 activity. Notably, previous studies have reported that TORC1-dependent Sch9 phosphorylation is inhibited by mitochondrial depolarization (64), coupling mitochondrial OXPHOS activity with cellular nutrient signaling. Since mitochondrial membrane potential is reduced in *ups1*Δ cells grown in glucose media (33), it is conceivable that mitochondrial depolarization upon inhibition of PA transport interferes with TORC1 signaling in *ups1*Δ cells. While further studies are needed to elucidate the link between mitochondrial depolarization and TORC1 signaling, our findings highlight that disturbances in the intramitochondrial transport of phospholipids and alterations in the phospholipid composition of mitochondrial membranes can broadly affect cellular signaling.

### Experimental procedures

#### Yeast strainsa

*Saccharomyces cerevisiae* strains used in this study are based on S288c or W303. The genotype of the yeast strains is described in the strain list table. Genes were deleted by PCR-targeted homologous recombination with the respective forward and reverse primers. Plasmids encoding disruption maker, HphNT1, NatNT2 or KanMX6, were used as template. C-terminal 6xHA tag of Sch9 at its endogenous loci was constructed using PCR-targeted homologous recombination with plasmid encoding 6HA-NatNT2 (65). For construction of cells expressing both Ire1(ΔIII)-3HA-GFP and *4xUPRE-GFP*, *4xUPRE-GFP* was integrated by PCR-targeted homologous recombination at the *URA3* locus in cells expressing Ire1(ΔIII)-3HA-GFP kindly provided by R. Ernst (PZMS, Homburg). *4xUPRE-GFP-URA3* was amplified using as a template genomic DNA of a yeast strain, which contained this gene and which was kindly provided by M. Graef (MPI ageing, Cologne) as a template.

#### Growth conditions

Yeast strains were cultured in YPD medium (1% yeast extract, 2% peptone, and 2% glucose) or synthetic complete glucose (SCD) medium (0.17% yeast nitrogen base without amino acids, ammonium sulfate, 0.5% ammonium sulfate, and 2% glucose) supplemented with necessary amino acids at 30°C. Choline (1 mM; SIGMA) in water, tunicamycin (0.25, 0.5 or 1 μg/ml; Merck) in DMSO or rapamycin (0.2 μg/ml; SIGMA) in DMSO were added to the media when indicated. For cell growth assay, cells were grown in SCD plate for 1 day at 30°C, and suspended in water at a concentration of 0.1 OD_600_ units. 4 μL of 5-fold serial dilutions of cells were spotted on SCD plates and incubated for 2days at 30°C.

#### Preparation of ER-enriched microsomes

100 OD_600_ units of cells grown to log phase in SCD medium were collected by centrifugation (3,000 × g, 5 min), washed once with H2O, resuspended in Tris-DTT buffer (0.1 M Tris, 10 mM DTT), and incubated for 15 min at 30 °C. Cells were collected by centrifugation (3,000 × g, 5 min), washed once with 1.2 M sorbitol, resuspended in sorbitol-phosphate buffer (20 mM potassium phosphate buffer [pH 7.4], 1.2 M sorbitol) containing lyticase, and incubated for 40 min at 30 °C. Spheroplasts were collected by centrifugation (3,000 × g, 5 min) and resuspended in ice-cold homogenization buffer (0.6 M sorbitol, 10 mM Tris-HCl [pH 7.4], 1 mM EDTA, 0.3% BSA (fatty acid free), 1 mM PMSF). Whole cell homogenates were subjected to centrifugation (1,200 × g, 5 min, 4°C). Membrane and microsomes soluble fraction were separated by centrifugation (17,500 × g, 5 min, 4°C). Supernatant was transferred to new tubes and subjected to centrifugation (40,000 × g, 40 min, 4°C). The pellet was resuspended in 200 μl of pure water and the protein concentration was determined by BCA assay. Aliquots of 50 μg protein were kept at −80°C and were subjected to lipid analysis by qMS.

#### Quantitative mass spectrometry of phospholipids

0.5 OD_600_ unit of cells or 5 μg protein of ER-enriched microsomes were used for lipid analysis by qMS.

Mass-spectrometric analysis of phospholipids was performed essentially as described (66). Briefly, lipids were extracted from samples in the presence of internal standards of major phospholipids (PC 17:0-14:1, PE 17:0-14:1, PI 17:0-14:1, PS 17:0-14:1, PA 17:0-14:1, PG 17:0-14:1 all from Avanti Polar Lipids) and CL (CL mix I, Avanti Polar Lipids LM-6003). Extraction was performed according to Bligh and Dyer with modifications. Final lipid samples were dissolved in 10 mM ammonium acetate in methanol and were sprayed into a QTRAP 6500 triple quadrupole mass spectrometer (SCIEX) by nano-infusion spray device (TriVersa NanoMate with ESI-Chip type A, Advion). The quadrupoles Q1 and Q3 were operated at unit resolution. Phospholipid analysis was carried out in positive ion mode. PC analysis was carried out by scanning for precursors of m/z 184 at a collision energy (CE) of 37 eV. PE, PI, PS, PG and PA measurements were performed by scanning for neutral losses of 141, 277, 185, 189 and 115 Da at CEs of 30, 30, 30, 25 and 25 eV, respectively. CL species were identified by scanning for precursors of the masses (m/z 465.4, 467.4, 491.4, 493.4, 495.4, 505.5, 519.5, 521.5, 523.5, 535.5, 547.5, 549.5, 551.5, 573.5, 575.5, 577.5, 579.5, 601.5, 603.5, 605.5, 607.5, 631.5, 715.5 and 771.5 Da) corresponding DAG-H2O fragments as singly charged ions at CEs of 45 eV. Mass spectra were analysed by the LipidView Software Version 1.2 (SCIEX) for identification, correction of isotopic overlap and quantification of lipids. Correction of isotopic overlap in CL species was performed using a spreadsheet calculating and subtracting theoretical amounts of [M+2] and [M+4] isotopes of each CL species that are isobalic to other CL species. Lipid amounts (pmol) were corrected for response differences between internal standards and endogenous lipids.

#### Immunoblotting

1 OD_600_ units of cells were collected, washed with H_2_O, and extracted by alkaline lysis (0.24 M NaOH, 1% 2-mercaptoethanol, 1 mM PMSF). Protein was precipitated in 25% trichloroacetic acid for 10 min on ice. Precipitates were washed with ice-cold acetone two times, dried, resuspended in 50 μL protein sample buffer (60 mM Tris-HCl [pH 6.8], 2% SDS, 10% glycerol, 20 mM DTT, 0.025% bromophenol blue), and incubated for 20 min at 40°C. Samples corresponding to 0.2 OD_600_ units of cells were separated by SDS-PAGE, immunobloted to nitrocellulose membrane, and immunodecorated with α-GFP (1:5000, ORIGENE), α-PGK1 (1:10,000, Abcam), or α-HA (1:1,000, Roche). Anti-rabbit, anti-mouse or anti-rat secondary antibodies conjugated to Dylight 800 (1:40,000, LI-COR) were used and detected with the Odyssey Infrared Imaging System (LI-COR). Quantification of the signals was performed using Image Studio Lite (LI-COR).

#### RNA extraction and quantitative RT-PCR

Total RNA was isolated with the NucleoSpin® RNA kit (MACHEREY-NAGEL) from 3 OD_600_ units of cells, according to the manufacturer’s protocol. cDNA was synthesized with 1 μg RNA and the GoScript™ Reverse Transcription Mix, Oligo(dT) (Promega). Quantitative RT-PCR was performed with Power SYBR Green PCR Master Mix (Thermo Fisher Scientific) and the following primers: *ACT1* forward 5’-TGTCACCAACTGGGACGATA and reverse 5’-AACCAGCGTAAATTGGAACG; spliced *HAC1* forward 5’-GCGTAATCCAGAAGCGCAGT and reverse 5’-GTGATGAAGAAATCATTCAATTCAAATG. QuantStudio 5 (Thermo Fisher Scientific) was used for measurement. Data were analyzed according to the ΔΔCT method normalized to housekeeping gene *ACT1*.

#### RNA sequence for transcriptomics

Libraries were prepared using the Illumina® TruSeq® mRNA stranded sample preparation Kit. Library preparation started with 1 μg total RNA. After poly-A selection (using poly-T oligo-attached magnetic beads), mRNA was purified and fragmented using divalent cations under elevated temperature. The RNA fragments underwent reverse transcription using random primers. This is followed by second strand cDNA synthesis with DNA Polymerase I and RNase H. After end repair and A-tailing, indexing adapters were ligated. The products were then purified and amplified (14 PCR cycles) to create the final cDNA libraries. After library validation and quantification (Agilent tape station), equimolar amounts of library were pooled. The pool was quantified by using the Peqlab KAPA Library Quantification Kit and the Applied Biosystems 7900HT Sequence Detection System. The pool was sequenced with a PE100 run on a NovaSeq6000 sequencer.

#### Bioinformatics and data analysis for transcriptomics

Raw reads were quantified using the alignment free quantification tool kallisto version 0.45.0. Reference genome is sacCer3. Gene counts were imported to R version 3.5.1 and normalized to library size using DESeq2 version 1.22.2. General differential gene expression was determined in pairwise comparisons using DESeq2. Functional enrichment of differentially expressed genes was performed using the DAVID API as previously reported (67). The Yeast GO-slim list was obtained from the *Saccharomyces* Genome Database.

#### Protein lysis and digest for proteomics

1 OD_600_ units of cells were lysed in 40 μL of 2% SDC in 100 mM Tris-HCl [pH 8.0] and lysate was cleared by centrifugation (12,000 rpm, 10 min, 25°C). Supernatant was subjected to protein concentration determination. In total 30 μg of protein were used for protein digestion. Proteins were reduced and alkylated by TCEP (10 mM) and CAA (20 mM) for 60 min at 45°C. 1 μg of LysC endopeptidase was added and incubated at 37°C or 2h followed by addition of 1 μg of Trypsin for digestion at 37°C for 16h. Digestion was stopped by addition of TFA to a final concentration of 0.5%. Lysates were cleared (SDC precipitates) by centrifugation and the supernatant was subjected for desalting using the StageTip (material: SDB-RPS, Affinisep) technique (68).

To generate the peptide spectral library, 2 μL of each sample (all conditions pooled 1:1) was pooled and subjected to high pH reversed phase chromatography. The instrumentation consisted out of a ZirconiumTM Ultra HPLC and a PAL RTC autosampler. The buffer systems consisted out of two buffers. A) 10 mM ammonium hydroxide and B) 80% acetonitrile and 10 mM ammonium hydroxide. Peptides were separated according to their hydrophobicity using an in-house packed column (length = 40 cm, inner diameter = 200 μm, 2.7-μm beads, PoroShell, Agilent Technologies) column. The instrument communicated and were controlled using the software Chronos (Axel Semrau GmbH). The gradient length was 60 min and in total 12 fractions were collected (1/60 s) and subsequently concentrated using a SpeedVac to complete dryness. Peptides were dissolved in 10 μl 2% formic acid, 2.5% acetonitrile of which 3 μL were injected per LC-MS/MS run.

#### Liquid chromatography and tandem mass spectrometry (LC–MS/MS)

For the MS/MS spectra library generation, the QExactive HF-x operated in a Top22 data-dependent mode. MS1 spectra were acquired in a mass range of 350–1,750 m/z, using an AGC target of 3 × 106 and a resolution at 200 m/z of 60,000. MS/MS spectra were acquired at 15,000 resolution using an AGC target of 5 × 105 and a maximal injection time of 22 ms.

For DIA measurements, MS1 spectra were acquired using a resolution of 60,000 and an AGC target of 1 × 106. For MS/MS independent spectra acquisition, 48 windows were acquired at an isolation m/z range of 15 Th and the isolation windows overlapped by 1 Th. The isolation center range covered a mass range of 385–1,043 m/z. Fragmentation spectra were acquired at a resolution of 15,000 at 200 m/z using a maximal injection time of 22 ms and stepped normalized collision energies (NCE) of 24, 27, 30. The default charge state was set to 4.

#### Bioinformatics and data analysis for proteomics

The MS/MS data dependent spectra library was analyzed with MaxQuant 1.6.3.4 and the implemented Andromeda search engine (69) using default settings. The spectra were correlated against the Uniprot reference Yeast proteome (downloaded 06.2019). The output txt folder was then used to build the spectral library in Spectronaut. DIA runs were analyzed using default settings. The data were exported in pivot-table format and further processed in Perseus (70). For pairwise comparison a two-sided t-test was utilized. The FDR was calculated by a permutation-based approach using 500 permutations and a fudge factor s0 of 0.1 A protein was considered to be significantly differently expressed at a FDR of 5%. Not cutoff for the fold change was considered. 1D enrichments (71) were performed in Perseus software using the Gene Ontology (GO) annotations of the Uniprot database. The Yeast GO-slim list was obtained from the *Saccharomyces* Genome Database.

#### SCH9 phosphorylation assay

2-nitro-5-thiocyanobenzoic acid (NTCB) treatments were performed as previously reported with slight modifications (54). 10 OD_600_ units of cells were treated with 6% trichloroacetic acid for 5 minutes on ice, washed twice with ice-cold acetone, and dried using a SpeedVac. The pellets were re-dissolved in 150 μl of urea lysis buffer (50 mM Tris [pH 7.5], 5 mM EDTA, 6 M urea, 1% SDS, 1 mM PMSF, and 1× Protease/Phosphatase inhibitor cocktail (CST)) and lysed with 0.7-mm-diameter zirconia beads in beads beater (3 × 30 s). After incubation for 10 min at 65°C and centrifugation (20,000 × g, 2 min), 100 μl of supernatant was transferred to a new 1.5 ml reaction tube. The lysates were mixed with 30 μl of 0.5 M CHES [pH 10.5] and 20 μl of 7.5 M NTCB and incubated overnight at room temperature. Each sample was mixed with 50 μl of 4× protein sample buffer (240 mM Tris-HCl [pH 6.8], 8% SDS, 40% glycerol, 80 mM DTT, 0.1% bromophenol blue) and subjected to western blotting.

#### Statistical analysis

Quantitative data are presented as arithmetic means ± standard deviation (SD). The statistical significance in related figures was assessed using two-tailed Student’s t test. *P* values and the number of experiments (n) are described in corresponding figure legend.

## Data and materials availability

All materials are available from the corresponding author upon request. Proteomics data will be deposited to the ProteomeXchange Consortium via the PRIDE partner repository. Transcriptomic data will be deposited to the GEO omnibus.

## Acknowledgments

We are grateful to Martin Graef, Robert Ernst, and Robbie Loewith for the kind gifts of plasmids and yeast strains. We thank the Cologne Center for Genomics for performing RNA sequencing and the MPI ageing bioinformatics facility for support in data processing. We also thank other lab members for discussion.

## Author contributions

A.E., T.T. and T.L. conceived and designed the research. A.E. performed the majority of experiments and data analysis. M.J.A generated strains and conducted initial analysis and growth assays. H.N. performed proteomic analysis. A.E., T.T. and T.L. wrote the manuscript.

## Funding and additional information

This work was supported by JSPS Overseas Research Fellowships to A.E. and by grants of the Deutsche Forschungsgemeinschaft to T.T. (TA1132/2-1) and T.L. (LA918/14-1).

